# Spatial updating and cue combination under simulated central visual field loss in virtual reality

**DOI:** 10.64898/2026.11.25.769221

**Authors:** Maggie K. McCracken, Maisha Tahsin Orthy, Sam Beech, Bobby Bodenheimer, Jeanine K. Stefanucci, Sarah H. Creem-Regehr

**Affiliations:** Department of Psychology, University of Utah; Department of Health & Kinesiology, University of Utahsss; Department of Computer Science, Vanderbilt University

**Keywords:** Spatial Updating, Cue Combination, Central Vision Loss, Virtual Reality

## Abstract

Central visual field loss can impair the use of visual landmarks for navigation, but whether it also changes how landmark and body-based cues are combined to support spatial updating remains poorly understood. We examined how gaze-contingent simulated central vision loss influenced variability, cue combination, and accuracy during spatial updating in virtual reality. Participants completed a homing task, in which they walked along two legs of a triangle and returned to their home location, under typical vision and with a simulated central scotoma in four cue conditions: landmark only, body-based only, both cues presented together, and cues presented in spatial conflict. Variability was generally reduced when both cues were available compared to single cue conditions, replicating the expected multisensory benefit, but similarly for both the scotoma and typical vision conditions. Participants consistently weighted visual landmarks more heavily than predicted by maximum likelihood estimation, and the simulated scotoma did not alter this weighting strategy. In contrast, simulated central vision loss selectively increased homing error when visual landmarks were available. Together, these findings suggest that central vision primarily supports accurate landmark processing during spatial updating, indicating that visual impairment degrades the quality of landmark information without fundamentally altering multisensory integration.

---

Successful navigation depends on the ability to continuously update one’s position while moving through an environment, a process called spatial updating (Klatzky et al., 1998; Wolbers et al., 2008). During spatial updating, navigators rely on both visual landmarks and body-based self-motion cues to estimate their position (Newman et al., 2023). Because visual landmarks often provide highly reliable information about location and orientation (Chan et al., 2012; Chen et al., 2017; Newman & McNamara, 2022), reducing the availability of landmark information may alter how navigators combine visual and body-based cues. Visual field loss, particularly loss of the central visual field, may reduce the reliability of visual landmark information due to reduced-acuity vision (blur) or the inability to accurately perceive cues for distance or location where an observer is fixating (Legge et al., 2016). However, it remains unclear whether this degradation of landmark information changes how navigators combine cues during spatial updating. Recent advances in immersive virtual reality (VR) and gaze-contingent eye tracking make it possible to simulate central visual field loss while maintaining precise control over both the visual environment and the characteristics of the simulated visual impairment. The present study uses this approach to examine how simulated central visual field loss influences cue combination during spatial updating.

## Spatial Updating and Cue Combination

Spatial updating relies on the continuous tracking of one’s position as one moves through space (Loomis et al., 1998). This process is supported by two primary sources of information: body-based cues, including vestibular and proprioceptive information about head motion and body movement, and external landmarks that provide information about relative position and orientation (Chen et al., 2017; Newman et al., 2023; Zhao & Warren, 2015a). Although landmarks can be perceived through multiple sensory modalities, including auditory, haptic, and olfactory (Schwarz & Hamburger, 2022), the present study focuses specifically on visual landmarks.

Because visual landmark and body-based information are inherently uncertain, the reliability of each source determines how much it should contribute to estimating one’s position during spatial updating. Combining information from multiple sources allows navigators to improve the precision of their position estimates, with each cue weighted according to its reliability (Ernst & Banks, 2002; Nardini et al., 2008). Navigators often exhibit a multisensory benefit, with reduced error and variability when both visual and body-based information are available, compared to either source alone (Chen et al., 2017; Newman & McNamara, 2021; Shayman, McCracken, Finney, Fino, et al., 2024). Maximum likelihood estimation (MLE) provides a normative framework for combining cues based on their relative reliability (Ernst & Banks, 2002). Under MLE, each cue is weighted according to its reliability (the inverse of its variance), such that more reliable cues contribute more to the final estimate. Consequently, MLE predicts both the relative weight assigned to each cue and that the combined estimate should be more precise than estimates based on either cue alone (Ernst & Banks, 2002; Nardini et al., 2008). Cue weights are commonly estimated using a spatial conflict paradigm, in which visual and body-based cues specify different locations, allowing the relative influence of each cue on navigators’ responses to be compared with MLE predictions (Newman et al., 2023). Using this approach, numerous studies have demonstrated MLE-consistent cue integration during spatial updating under typical viewing conditions (Chen et al., 2017; Nardini et al., 2008; Newman et al., 2023; Shayman, McCracken, Finney, Katsanevas, et al., 2024).

Although MLE provides a normative framework for cue combination, researchers have shown that cue weighting during spatial updating changes systematically as cue reliability is manipulated (Chen et al., 2017; Zanchi et al., 2022; Zhao & Warren, 2018). For example, navigators assign greater weight to landmark information when landmarks are more numerous or located closer to the home location, thereby increasing their reliability (Chen et al., 2017; Zanchi et al., 2022; Zhao & Warren, 2015a). Conversely, reducing the reliability of a cue leads navigators to down-weight that cue and rely more heavily on alternative sources of information (Roy et al., 2023; Zhao & Warren, 2015a). When differences in cue reliability become sufficiently large, navigators may abandon cue integration altogether and rely exclusively on the more reliable cue (Zhao & Warren, 2015a, 2018). Observers may also integrate cues in ways that are inconsistent with a cue’s reliability (Chrastil et al., 2019; Roy et al., 2023; Zhang et al., 2020).

Much of the previous spatial updating work has been conducted using VR homing paradigms, in which participants walk along a path to one or more locations before attempting to return to a remembered home location (Chen et al., 2017; Newman et al., 2023; Riecke et al., 2002; Xie et al., 2017). During these tasks, visual landmarks, or stable, localizable objects in the environment (e.g., signs, trees, or small buildings), provide external spatial references, while vestibular and proprioceptive cues generated during natural walking provide body-based information about self-motion. By allowing participants to walk naturally, ambulatory immersive VR preserves these body-based cues while providing precise control over visual landmarks, making it well suited for studying cue combination during spatial updating (Klatzky et al., 1998). Furthermore, cue combination paradigms with conflict conditions can quantify the benefits of combining cues, estimate cue weighting, and compare observed cue combination with predictions from MLE. Comparing single-cue conditions, in which only visual or body-based information is available, with conditions in which both cues are available allows researchers to determine the contribution of each cue and whether combining them reduces variability relative to either cue alone. In conflict conditions, the cues provide different estimates of the remembered home location, allowing the relative influence of each cue to be quantified from the participant’s response. This approach is particularly useful to simulate changes in sensory capabilities that are associated with different populations – such as older adults or people with visual impairment – to understand subsequent multisensory integration effects on navigation performance.

## Central Vision Loss and Navigation

Visual impairment provides a natural test of how changes in cue reliability influence spatial updating. Vision loss is highly prevalent, affecting approximately 441.5 million individuals worldwide whose vision cannot be fully corrected, defined as low vision (Bourne et al., 2021). Low vision encompasses a range of visual impairments, including reduced acuity, reduced contrast sensitivity, and visual field loss, each of which can differentially affect spatial behavior (Creem-Regehr et al., 2021). Despite reduced visual input, basic spatial updating performance is often relatively preserved across a range of visually degraded conditions because body-based information can effectively support self-localization (Legge et al., 2016; Loomis et al., 1993; Tarampi et al., 2010) and/or high-contrast object boundaries can still be used (Freedman et al., 2019; Rand et al., 2012; Shayman, McCracken, Finney, Fino, et al., 2024). The comparable performance in spatial updating and other mobility and navigation tasks may be due to adopting compensatory behaviors, including increased visual scanning through head movements and greater reliance on nonvisual sources of spatial information (Cloutier & DeLucia, 2022).

Although non-visual information can support spatial updating when vision is degraded, not all forms of vision loss are expected to have equivalent consequences for performance. Central visual field loss may represent a particularly important case because a central scotoma degrades the high-resolution foveal information used to identify and localize visual landmarks while leaving peripheral vision largely intact. As a result, navigators must rely on lower-acuity peripheral vision to obtain landmark information, potentially reducing the reliability of visual landmarks as a spatial cue. Our prior work manipulated visual cue reliability using acute simulated blur (20/400) across the full visual field in virtual reality, which did not alter visual and auditory cue weighting or increase homing variability during spatial updating (Shayman, McCracken, Finney, Fino, et al., 2024). A central vision scotoma that is yoked to eye movements may have different consequences because it selectively blocks the area surrounding fixation, prohibiting direct viewing of the landmarks. A recent study with simulated central scotomas showed reduced navigation efficiency, increased the time required to detect visual targets, and altered gaze behavior during naturalistic search, suggesting that eye-tracked scotomas may substantially degrade the quality of visual information available to support navigation (Veerkamp et al., 2025).

Our understanding of the influence of central vision loss on mobility and navigation comes from both individuals with clinically-defined vision loss (e.g., age-related macular degeneration, AMD) and simulations implemented on typically sighted individuals. Individuals with clinical central vision loss have shown slower walking speeds and reduced accuracy in foot placement when tasked with walking in low light, suggesting that reduced visual information influences mobility associated with navigation (Alexander et al., 2014a, 2014b; Napier-Dovorany et al., 2026). Differences in gaze patterns and head movements are also identified in people with AMD when navigating to a location (Hassan et al., 2002) or making decisions about street crossing (Geruschat et al., 2006; Hassan et al., 2005). Many prior simulations of lower acuity central vision functioning in real world spaces have used blur-filter goggles. These studies have shown that degraded acuity and contrast sensitivity affect the accuracy of recognition of environmental features important for safe locomotion (Bochsler et al., 2012), overestimation of perception of speed of locomotion (Rand et al., 2019), perceived locomotion speed (Rand et al., 2019), and impaired spatial memory for landmarks encountered during navigation (Barhorst-Cates et al., 2017; Rand et al., 2015). These spatial learning and memory studies identified higher cognitive load associated with the increased uncertainty of navigating under degraded vision conditions (Barhorst-Cates et al., 2017).

While informative, the real world simulations face limitations, including a lack of ability to isolate the consequences of specific visual field loss. More recently, eye-tracking capabilities in head-mounted displays and virtual environments enable the rendering of more precise visual scotomas tied to one’s gaze. These have been used to simulate eye-centered deficits such as cataracts (Krösl et al., 2020) to examine their effects on daily tasks such as reading signs (Krösl et al., 2018) as well as to provide experiences for sighted individuals that could improve empathy and understanding (Krösl et al., 2023). This gaze-contingent method was used in Orthy et al. (2026), showing that simulated gaze-contingent peripheral field loss affected observers’ judgments of their abilities to step over a virtual gap. Taken together, prior work establishes the importance of central vision for perceiving and locating environmental features such as landmarks, and shows that, in some cases, tasks can be completed as successfully with visual impairment as with typical vision. However, the mechanisms underlying navigation with visual impairment are not well-understood (Cloutier & DeLucia, 2022). Our approach to examine how multisensory cues are integrated with simulated central visual field loss begins to address this gap in understanding. There is some suggestion that multisensory (audiovisual) integration did not follow MLE predictions in a spatial localization task with people with progressive central vision loss (Garcia et al., 2017), but there has been no investigation of multisensory cue integration for navigation in the context of selective vision loss.

## The Current Study

The current study examined whether simulated central visual field loss alters cue combination during spatial updating. Central visual field loss was simulated using a gaze-contingent scotoma that selectively restricted central visual information while preserving peripheral vision and body-based self-motion cues. Participants completed an ambulatory virtual reality homing task under both typical vision and simulated central visual field loss. Spatial updating was assessed using three complementary measures: homing variability, cue weighting relative to predictions from maximum likelihood estimation (MLE), and homing error.

Because central visual field loss is expected to reduce the reliability of visual land-mark information, we examined three primary hypotheses. First, we predicted a smaller multisensory benefit with simulated central vision loss than with typical vision, because the landmark information would be less reliable (**H1**). This would lead to reduced variability under consistent cues compared to single-cue conditions with typical vision, but no (or a smaller) advantage of consistent cues over single-cue conditions when viewing with the scotoma. In addition to changes in variability, we predicted that integration processes would change as well. Specifically, we predicted that multisensory variability would match MLE predictions under typical vision but would be greater than predicted under simulated scotoma, indicating a reduced multisensory benefit (**H2a**). Additionally, we predicted that under typical vision, cue weights would be weighted towards vision and consistent with MLE predictions, whereas the simulated scotoma would result in greater weighting of body-based cues relative to visual landmarks (**H2b**). Third, we predicted that reduced access to landmark information would increase homing error (**H3**). This would lead to greater error in the conditions that allow for landmark use during homing (consistent and landmark) in the presence of a scotoma compared to typical vision.

## Method

### Participants

Thirty-two undergraduate students (*M_age_* = 20.7; 17 female, 15 male) were recruited from the University of Utah participant pool in exchange for course credit. The study protocol was approved by the university’s Institutional Review Board, and all participants gave written, informed consent to participate. The sample size was determined via an *a priori* power analysis with a medium effect size (*η*^2^ = 0.11-0.18) reported in previous VR homing task studies (Chen et al., 2017; Newman & McNamara, 2022; Sjolund et al., 2018). We found that 26 participants would be sufficient to find within-group effects at 80% power in a general linear model. We over sampled this number given potential technical failures of the virtual reality system.

All participants had at least 20/40 or corrected-to-20/40 vision verified using a Snellen chart, reported no history of neurological or visual disorders, and were able to walk without assistance. To screen for balance abnormalities, participants completed a Romberg test in which they stood on a foam pad with eyes closed for 30 seconds. Data from several participants were excluded from the analyses due to technical issues (*n* = 1) and experimenter procedural errors in the vision-only condition (*n* = 3), yielding a final sample of 28 participants.

### Materials

The virtual environment (VE) consisted of an endless grassy ground plane extending to a visible horizon; objects did not cast shadows. The VE was designed in Unity (build 2020.3.25f1) and rendered via the SteamVR Plugin (Version 2.7.3) and SteamVR (Version 2.13). A rendering of the VE is shown in Figure 1 (left). The VE was displayed using an HTC Vive Pro Eye head-mounted display (HMD) with a wireless adapter (resolution: 1440 × 1600 pixels per eye; refresh rate: 90 Hz; field of view: 110°). The experiment took place in a 8 × 9 m room with 6 × 6 m tracking space.

**Figure 1.**
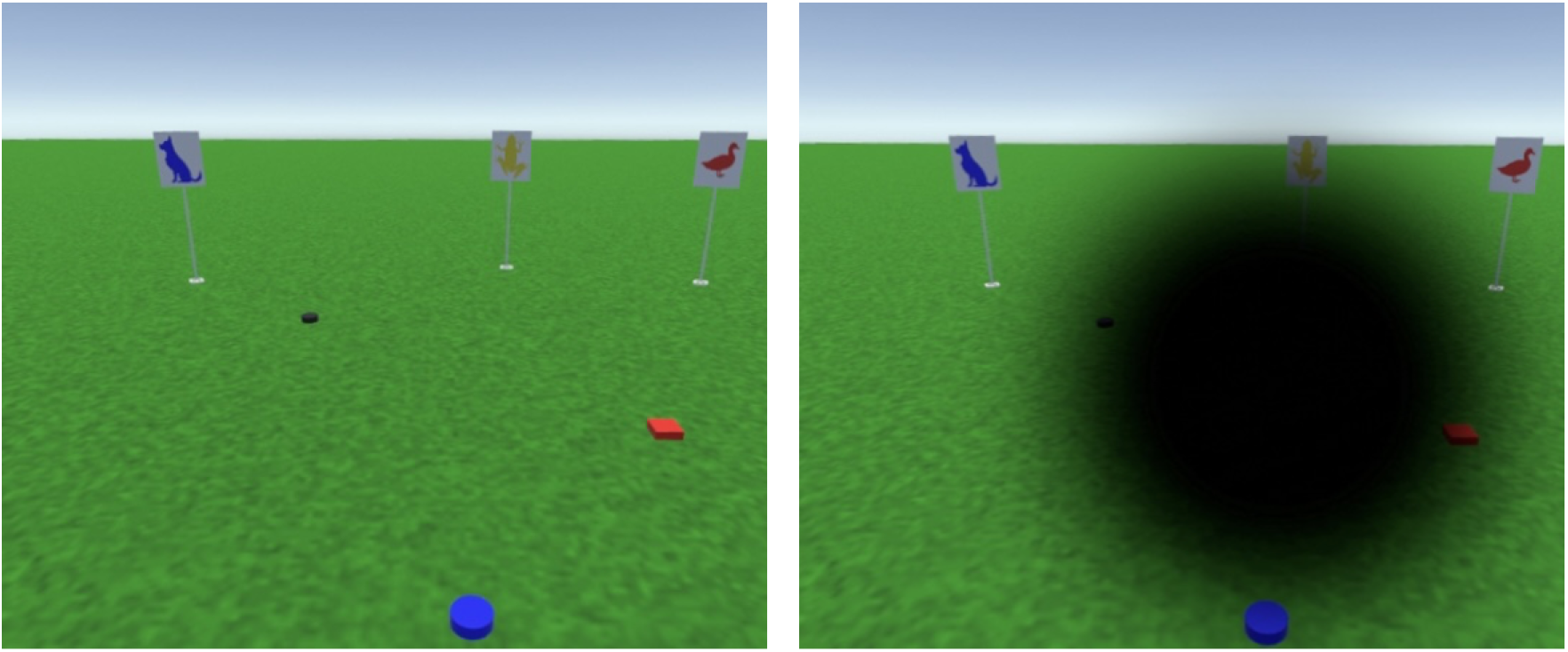
A rendering of the virtual environment (left) and a rendering of the simulated central scotoma (right)

Three visual landmarks with distinct figures were placed in the VE: a blue dog, a yellow frog, and a red duck. The task layout is illustrated in Figure 2 (left), with the second waypoint treated as the origin of the coordinate grid. Landmarks were positioned 4.5 m from the origin at −30*^◦^*, +7.5*^◦^*, and +30*^◦^* and were 1.45 m tall with poster dimensions of 36 cm × 50 cm.

**Figure 2.**
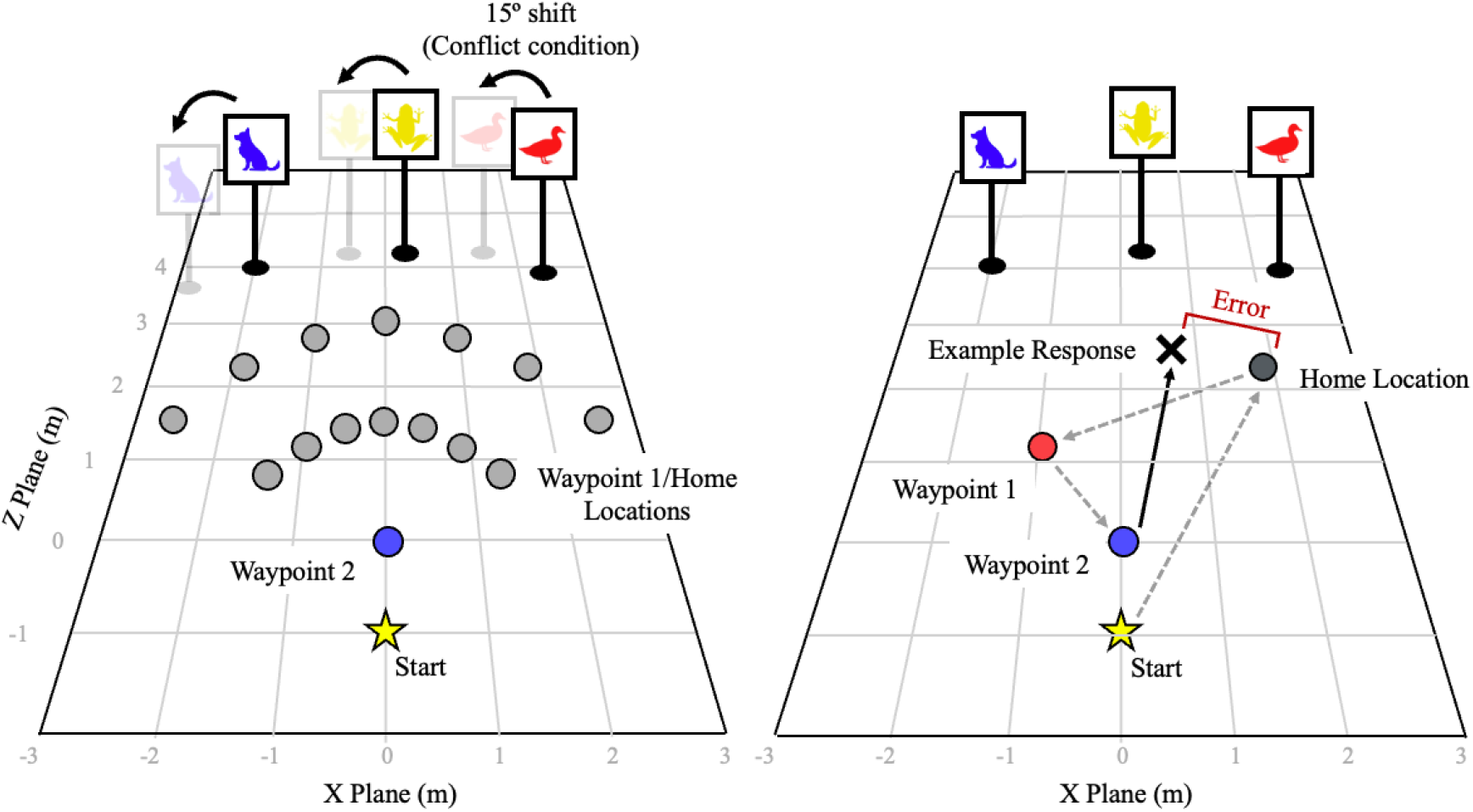
An overview of the experimental layout (left) and trial progression (right) from an overhead perspective.

Home and waypoint locations were denoted by three virtual discs (diameter: 12 cm, height: 4.75 cm): a black disc for the home location, a red disc for the first waypoint, and a blue disc for the second waypoint (grid origin). The home and waypoint locations were either 1.5 or 3 m from the grid origin. Eight unique triangular path configurations were used, each repeated once per condition, with two additional configurations reserved for practice trials. Total path lengths ranged from 1.4 to 4.0 m and angular positions spanned ±60° from the grid origin.

The scotoma was rendered as a gaze-centered, screen-space mask with a fixed pixel radius per eye. The angular extents were built using the manufacturer’s FOV (110*^◦^* diagonal) and a horizontal resolution of 1440 pixels, yielding a fully opaque core scotoma of approximately 28*^◦^* with a gradient boundary extending to approximately 53*^◦^* (Figure 1 right). The scotoma was intentionally large to produce a robust degradation of visual information and maximize potential effects on spatial updating, rather than to replicate a specific clinical pattern of visual field loss.

Throughout the homing task, participants wore in-ear wireless headphones (Beats Flex Wireless) playing continuous brown noise. Noise intensity was calibrated so that participants could hear verbal instructions from the experimenter but not ambient room sounds such as air conditioning or footsteps, minimizing the use of real-world auditory cues.

### Procedure & Design

The experiment used a 2 (vision condition: typical, central visual field loss) × 4 (cue condition: landmark only, body-based only, consistent, conflict) within-subjects design. After providing informed consent, participants completed Snellen visual acuity screening and a Romberg balance test. Participants closed their eyes and were guided into the laboratory before donning the HMD, ensuring they did not view the experimental room during the study visit. Eye tracking was then calibrated in the HMD using the HTC Vive Pro Eye’s built-in SRanipal eye-tracking system.

The homing task consisted of four phases on each trial: encoding, cue manipulation, response, and resetting. During the encoding phase, participants walked to the home location (black disc) and then to two subsequent waypoints (red and blue discs). Upon reaching the second waypoint, the participant was virtually blindfolded as the HMD screen turned black.

Participants remained virtually blindfolded for 10 seconds, during which one of four cue manipulations was applied. In the landmark-only condition, participants were gently spun in a swivel chair in an irregular sequence of left and right rotations, such that they could not track their final heading. In the body-based-only condition, landmarks were removed from the environment so that the response phase relied solely on body-based cues. In the consistent-cue condition, no manipulation occurred, and both landmarks and body-based cues remained available. In the cue-conflict condition, landmarks were covertly shifted 15° to the left around the origin during the blindfolded period, creating a spatial conflict between landmark and body-based cues during the response phase.

Following the blindfolded period, the HMD screen was restored, and participants completed the response phase by walking to the location they remembered as home with-out the discs. Response locations were recorded using head-tracked position data from the HMD. After the response, participants were blindfolded again and guided back to the starting location by the experimenter along an indirect path to minimize redundant distance information. An experimenter remained in close proximity to the participant throughout the experiment to ensure safety and prevent the participant from reaching the boundary of the tracking space.

The vision condition was a within-subject manipulation with participants randomly assigned to complete either the central visual field loss condition or the typical vision condition first, counterbalanced to prevent order effects. The scotoma was present throughout all trials in the vision loss condition and absent in the typical vision condition. Each vision condition consisted of 36 trials: the first four served as practice (one per cue condition), followed by 32 experimental trials with eight trials per cue condition presented in randomized order. Participants completed all trials in one vision condition before taking a 5-minute break, after which they completed the second vision condition. Upon completing both conditions, participants answered questions about their video game and VR experience. The total session lasted approximately 1.5 hours.

### Data Analysis

All analyses were conducted in R (Version 4.6.0; R Core Team, 2026) using the *brms* package (Version 2.23.0; (Bürkner, 2021)), which fits Bayesian multilevel models via the probabilistic programming language Stan (Carpenter et al., 2017). Data and analysis scripts are publicly available on the Open Science Framework (OSF): https://osf.io/vjg4x/overview?view_only=427b2c4bd15445a8af8f7731ae31d195.

Homing error and variability were each modeled using Bayesian multilevel models with cue condition, vision condition, and their interaction as fixed effects and a random intercept by participant to account for individual differences. Accuracy was operationalized as the Euclidean distance between the participant’s response location and the true home location on each trial (in cm), with larger values indicating greater homing error. Variability was calculated within each cue and vision condition as the standard deviation of the Euclidean distances from each trial’s response location to the participant’s centroid of responses. Weakly informative priors were specified based on previous work using the same homing paradigm (Shayman, McCracken, Finney, Fino, et al., 2024); complete prior specifications and prior predictive checks are provided in the Supplementary Materials. Convergence was assessed using *R̂* ≤ 1.01 and effective sample sizes greater than 1,000 for all parameters. All models converged successfully, with *R̂* = 1.00 and bulk ESS *>* 6000 for all parameters. All parameter estimates are reported as posterior means with 95% credible intervals (CrIs). Effects were considered credible when the 95% CrI excluded zero. Bayes factors (*BF*_10_) are reported as continuous measures of the relative evidence for the alternative versus the null hypothesis, with values greater than 1 favoring the alternative hypothesis and values less than 1 favoring the null hypothesis.

#### Cue Combination Modeling

Cue combination analyses compared predictions from an MLE model, derived from single-cue condition performance, to observed performance in the consistent and conflict conditions. The predicted weights (*w*^) for landmark (*l*) and body-based (*b*) cues were determined using the relative variance (*σ*^2^) of the single-cue conditions:

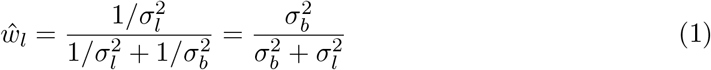

Because only two cues are available, *w*^*_l_*and *w*^*_b_* sum to unity:

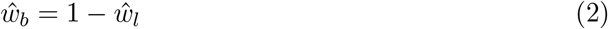

The predicted MLE variance for the consistent-cue condition is derived using the predicted weights and observed single-cue variances:

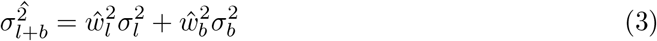

These model predictions are then compared to observed performance. Observed cue weights (*w*) are derived from the conflict condition, in which the 15^◦^ angular shift of the landmarks caused landmark and body-based cues to specify different home locations. First, response locations were transformed into a common coordinate system, with the second waypoint as the origin and the y-axis aligned with the correct walking direction toward the home location. This transformation ensures that the 15^◦^ shift of the landmarks in the conflicting-cue condition was in the same reference frame across the range of home locations. The distances between the centroid of responses in the conflict condition (*µ_conflict_*) and the centroids of the landmark-only and body-based-only conditions were calculated as:

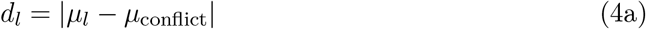

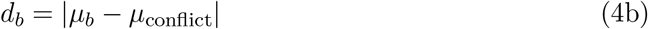

which is then used to compute the observed cue weighting that is compared to the MLE predictions:

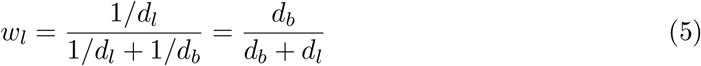

#### Head Rotation Processing

Head rotation during the homing leg was examined as an exploratory measure derived from the continuous HMD tracking data, recorded at 60 Hz. Yaw rotation rate (degrees per second) was calculated as the absolute rate of change in head yaw angle between consecutive frames in the horizontal plane only, reflecting left-right rotational movement of the head independent of changes in vertical (pitch) or lateral (roll) orientation. To exclude the initial turn used to orient toward the home location, analyses were restricted to frames in which participants were translating through the environment, defined by a frame-to-frame change in horizontal head position exceeding 0.5 cm. Yaw was computed by unwrapping the yaw signal to remove discontinuities at *±*180*^◦^* and dividing the frame-by-frame change in yaw by the corresponding change in time. Yaw rotation rate was averaged across frames within each trial, yielding one estimate per participant per trial. These trial-level estimates were then entered directly into the Bayesian multilevel model as the outcome variable.

## Results

### Variability & Cue Combination

We hypothesized a reduced multisensory benefit (i.e., a smaller advantage of combined cues over single cues) with central visual field loss due to reduced reliability of land-mark information (**H1**). To test this prediction, homing variability (standard deviation) was modeled using a Bayesian mixed model with cue condition, vision condition, and their interaction as fixed effects, a random intercept by participant, and the consistent-cue condition and typical vision condition were dummy coded as the reference groups (see Figure 3). To examine whether participants showed reduced variability (SD) in the consistent-cue condition relative to single-cue conditions under typical vision, we examined the main effects of cue condition. Baseline variability in the consistent-cues condition under typical vision was 29.88 cm (*B*_0_ = 29.88, 95% CrI [24.73, 34.77]). Variability was credibly greater in the landmark-only condition (*B* = 7.99, 95% CrI [0.68, 15.28], *BF*_10_ = 3.57). The body-based cues condition showed a similar pattern but with weaker evidence (*B* = 6.19, 95% CrI [*−*1.06, 13.53], *BF*_10_ = 1.92).

**Figure 3.**
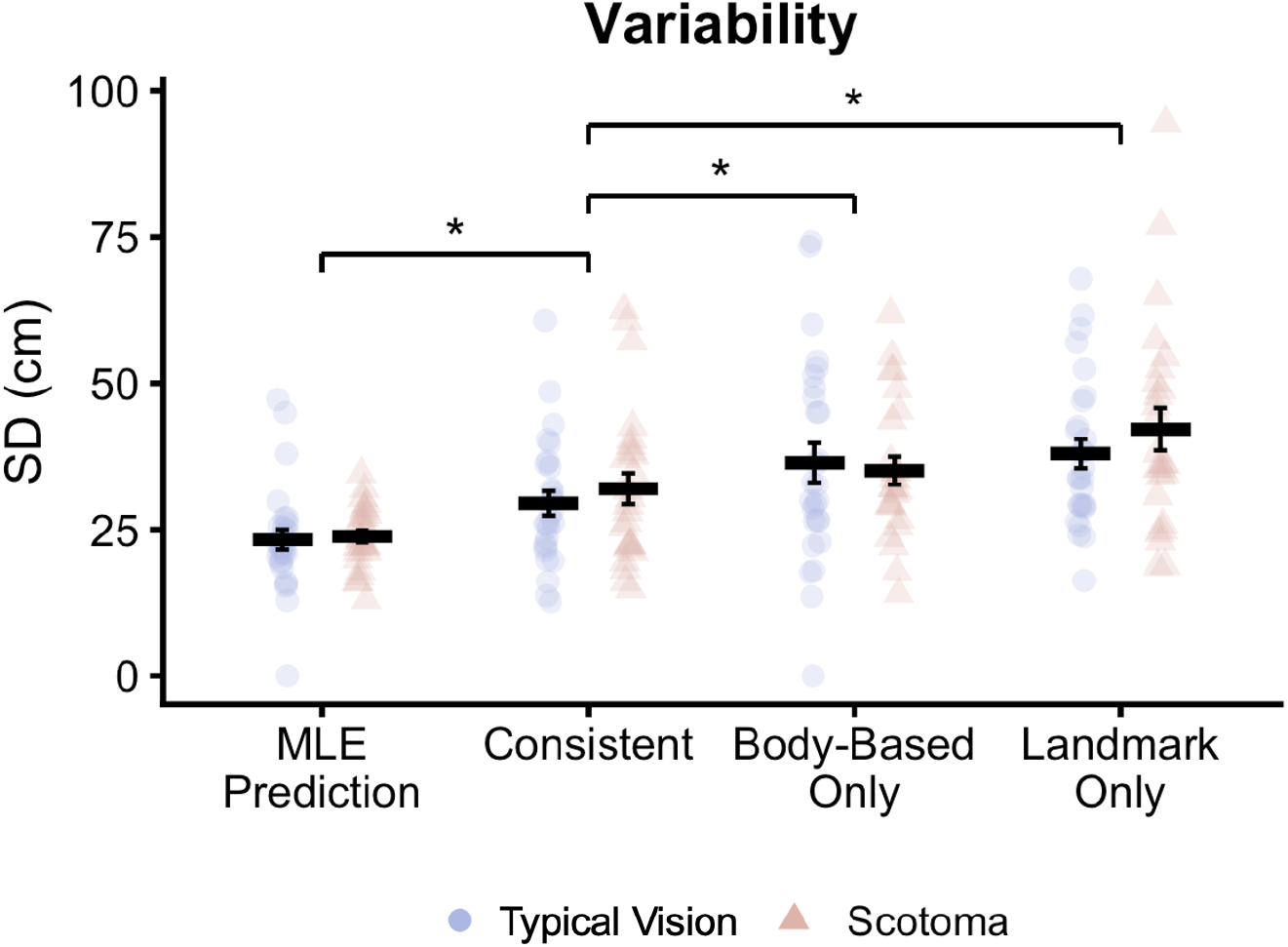
Homing variability (standard deviation) across cue conditions for typical vision (blue) and simulated scotoma (red). Points represent individual participant means, and black horizontal lines and error bars represent posterior means and 95% credible intervals, respectively, derived from Bayesian multilevel models. Asterisks indicate comparisons between single-cue and consistent-cue conditions or between observed consistent-cue variability and maximum likelihood estimation (MLE) predictions for which the 95% credible interval of the difference excludes zero.

To assess the multisensory benefit under simulated scotoma, we compared variability in the consistent-cues condition with that of each single-cue condition. Baseline standard deviation (SD) under simulated scotoma in the consistent-cues condition was 30.03 cm (*B*_0_ = 30.03, 95% CrI [24.90, 35.25]), comparable to typical vision. The landmark-only condition showed credibly greater SD than consistent cues (*B* = 7.79, 95% CrI [0.30, 15.03], *BF*_10_ = 2.13), though with weaker evidence than under typical vision. The body-based condition showed a similar pattern but with ambiguous evidence (*B* = 6.10, 95% CrI [*−*1.01, 13.38], *BF*_10_ = 0.96). The multisensory benefit for variability therefore showed a broadly similar pattern under simulated scotoma as under typical vision, with the clearest evidence of a benefit relative to landmark cues alone. Together, contrary to the predictions of **H1**, these results suggest that simulated central visual field loss did not substantially alter the pattern of multisensory benefits for homing variability across cue conditions, showing the expected reduction in variability given two consistent cues over single cues but for both viewing conditions.

To examine whether simulated central visual field loss affected homing variability, we compared the SD of homing responses between vision conditions. Homing variability did not credibly differ between typical vision and simulated scotoma in the consistent-cues condition (*B* = 2.05, 95% CrI [*−*5.02, 9.14], *BF*_10_ = 1.28), the landmark-only condition (*B* = 3.40, 95% CrI [*−*3.84, 10.73], *BF*_10_ = 1.35), or the body-based cues condition (*B* = *−*0.71, 95% CrI [*−*7.89, 6.52], *BF*_10_ = 1.27). Therefore, there was no credible evidence supporting a difference in the precision of homing performance in any cue condition.

To test whether combining consistent cues increased reliability as predicted by MLE (**H2a**), we compared observed homing variability (SD) under consistent cues with MLE-predicted variability derived from the single-cue conditions (see Figure 3). We modeled homing variability as a function of scotoma condition, variability type (predicted vs. observed), and their interaction, with a random intercept by participant. MLE-predicted SD under typical vision was 23.47 cm (*B*_0_ = 23.47, 95% CrI [19.84, 27.02]) and did not credibly differ under simulated scotoma (*B* = 0.42, 95% CrI [*−*4.26, 5.23], *BF*_10_ = 0.25). Observed SD was greater than predicted under typical vision (*B* = 5.79, 95% CrI [0.88, 10.45], *BF*_10_ = 3.45), indicating less multisensory benefit than predicted by MLE. This difference did not credibly change under simulated scotoma (*B* = 2.07, 95% CrI [*−*4.29, 8.71], *BF*_10_ = 0.41). Thus, contrary to **H2a**, multisensory benefit was smaller than predicted by MLE in both vision conditions.

To determine whether participants changed how they combined landmark and body-based information, and to examine whether simulated central visual field loss disrupted cue combination (**H2b**), we regressed vision cue weight onto scotoma condition, weighting type (predicted vs. observed), and their interaction, with a random intercept by participant (Figure 4). Under typical vision, predicted vision weights averaged 0.47 (*B*_0_ = 0.47, 95% CrI [0.38, 0.55]), indicating roughly equal weighting of visual and body-based cues as predicted by the MLE model. Observed vision weights were credibly higher than predicted (*B* = 0.14, 95% CrI [0.04, 0.25]), suggesting participants overweighted visual cues relative to MLE predictions under typical vision. Contrary to **H2b**, simulated central visual field loss did not credibly alter predicted weights (*B* = *−*0.02, 95% CrI [*−*0.13, 0.09]), as demonstrated with the single cue SD analyses above, nor did it credibly change the discrepancy between observed and predicted weights (interaction: *B* = *−*0.01, 95% CrI [*−*0.15, 0.14]). **H2b** was therefore not supported. While participants showed a systematic tendency to overweight visual cues relative to MLE predictions (shown via a relatively flat regression line above the 45*^◦^* parity line in Figure 4), this pattern did not differ credibly between typical vision and simulated scotoma conditions.

**Figure 4.**
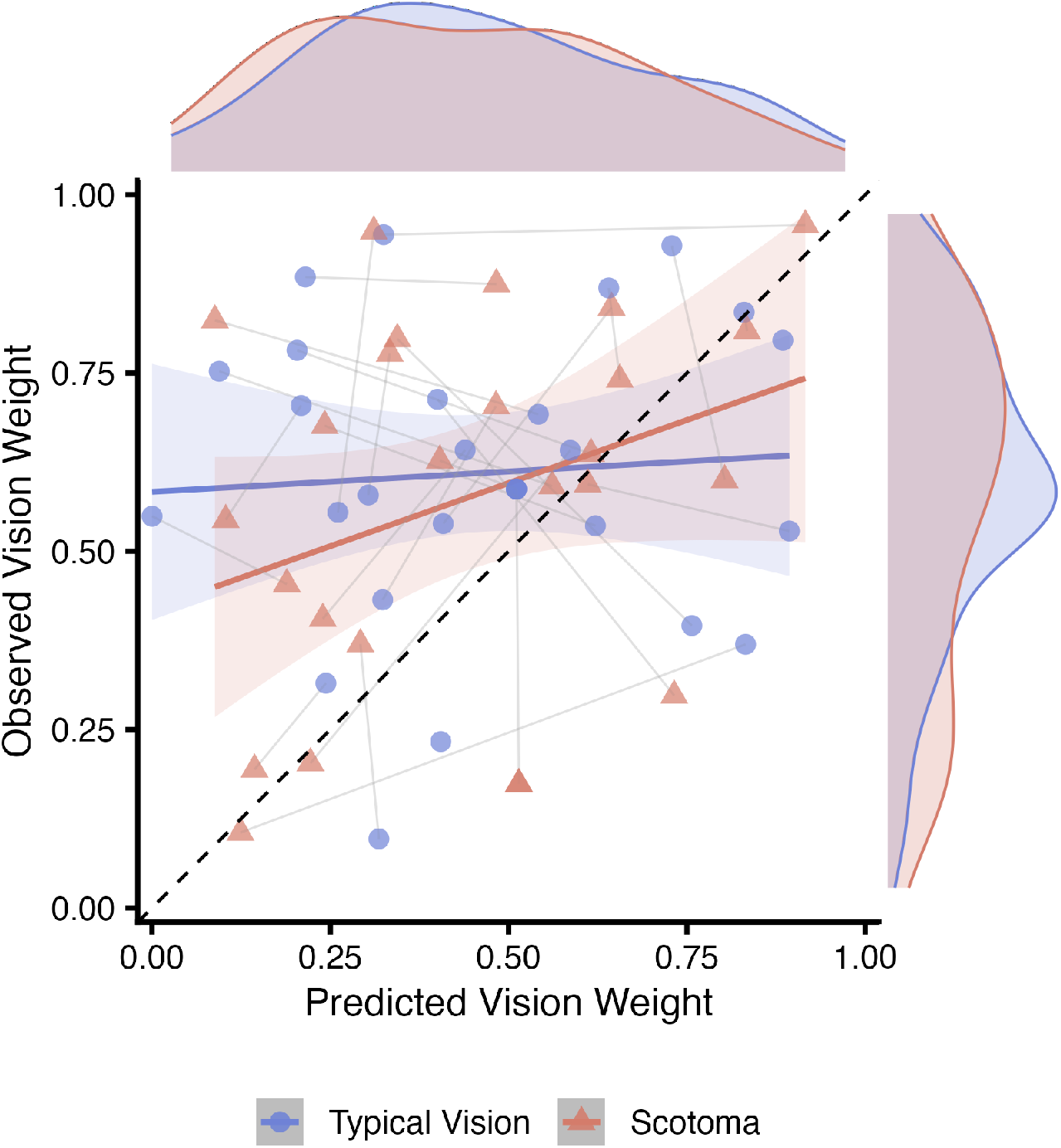
Observed versus predicted vision cue weights for each participant under typical vision (blue) and simulated scotoma (red). Each point represents one participant, with lines connecting the same participant across vision conditions. Marginal density plots show the distribution of predicted (top) and observed (right) weights by vision condition.

Because visual inspection of the data suggested variability in participants’ responses to the simulated central scotoma, we conducted an exploratory analysis by fitting a Bayesian multilevel model with a random slope for scotoma condition by participant (Figure 5). The standard deviation of the random slope for scotoma was small but non-zero (*SD* = 0.08, 95% CrI [0.00, 0.19]), indicating some between-participant variability in how simulated central visual field loss affected vision weighting. However, fixed-effects estimates were consistent with the primary model, given that the central scotoma did not credibly alter vision weights at the group level (*B* = *−*0.03, 95% CrI [*−*0.14, 0.08]). These findings suggest that some individuals adjusted their reliance on visual information under simulated central visual field loss, although these adjustments were not consistent enough to produce a reliable group-level effect.

**Figure 5.**
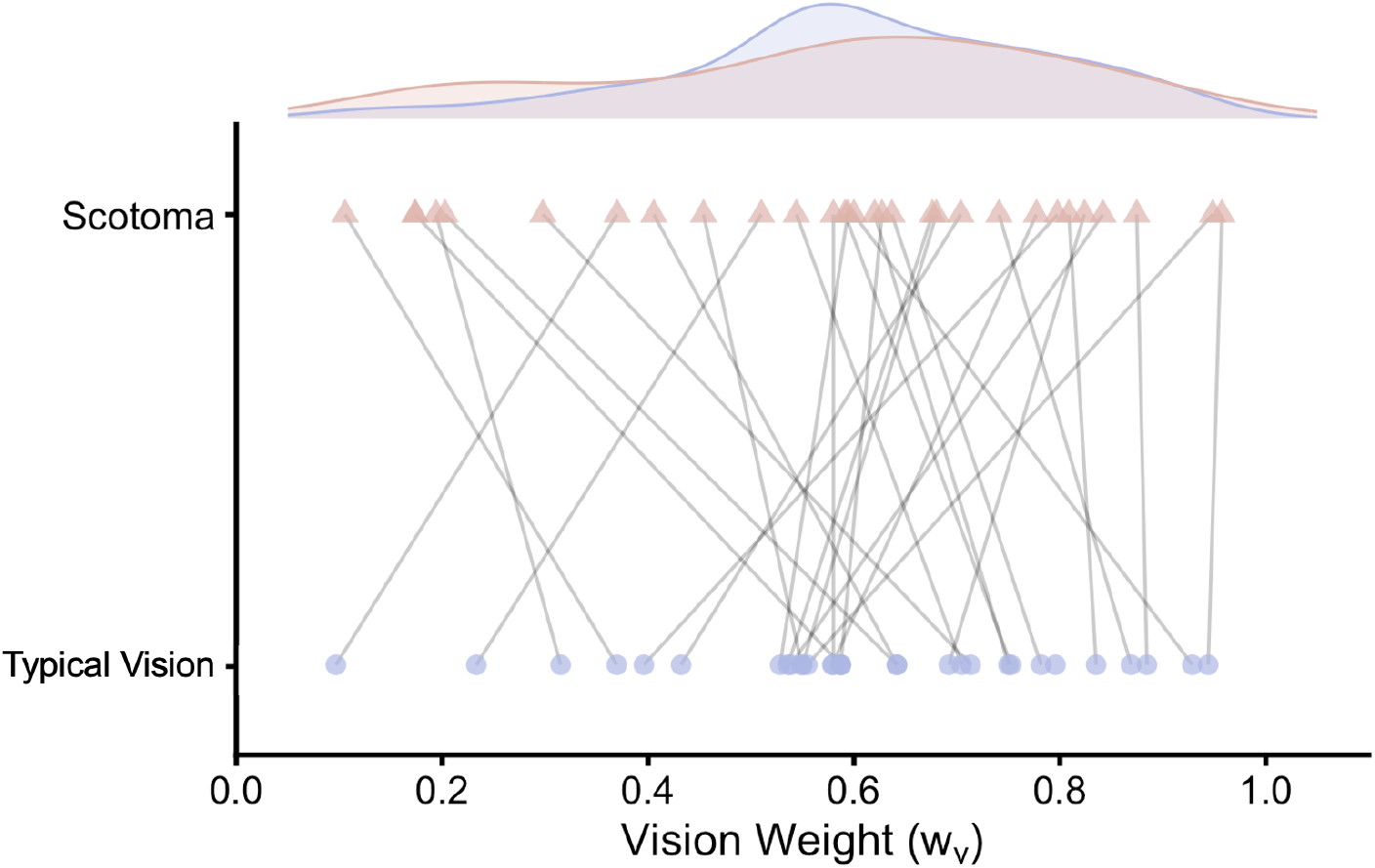
Visual weighting across viewing conditions. Observed visual weights (w_v_) for each participant under typical vision (blue) and simulated central scotoma (red). Lines connect each participant across conditions, and marginal density curves show the distribution of visual weights within each viewing condition.

### Accuracy

Our third hypothesis predicted that reduced access to landmark information would increase homing error (**H3**). Homing error was modeled on a trial-by-trial basis using a Bayesian mixed model with cue condition, scotoma condition, and their interaction as fixed effects and a random intercept by participant to account for individual differences (see Figure 6). There was weak evidence that homing error was greater under simulated scotoma than typical vision in the consistent-cues condition (*B* = 8.90, 95% CrI [0.28, 17.43], *BF*_10_ = 1.15). Homing error was credibly greater in the landmark-only condition (*B* = 11.53, 95% CrI [2.83, 20.23], *BF*_10_ = 6.25). Homing error in the body-based cues condition did not credibly differ between scotoma and typical vision (*B* = 2.15, 95% CrI [-6.32, 10.78], *BF*_10_ = 0.16).^1^ These results provide support for **H3**, with the strongest evidence for impaired homing accuracy when participants relied solely on visual landmarks and no evidence of impairment when participants relied on body-based cues alone. These findings indicate that the simulated central scotoma impaired navigation primarily when successful performance depended on visual landmarks. When body-based information was the primary cue, homing accuracy remained unaffected. Taken together with the SD results, which showed no difference between typical vision and simulated scotoma, these findings suggest that the scotoma introduced a systematic error rather than decreased precision (or increased variability) in navigation performance.

**Figure 6.**
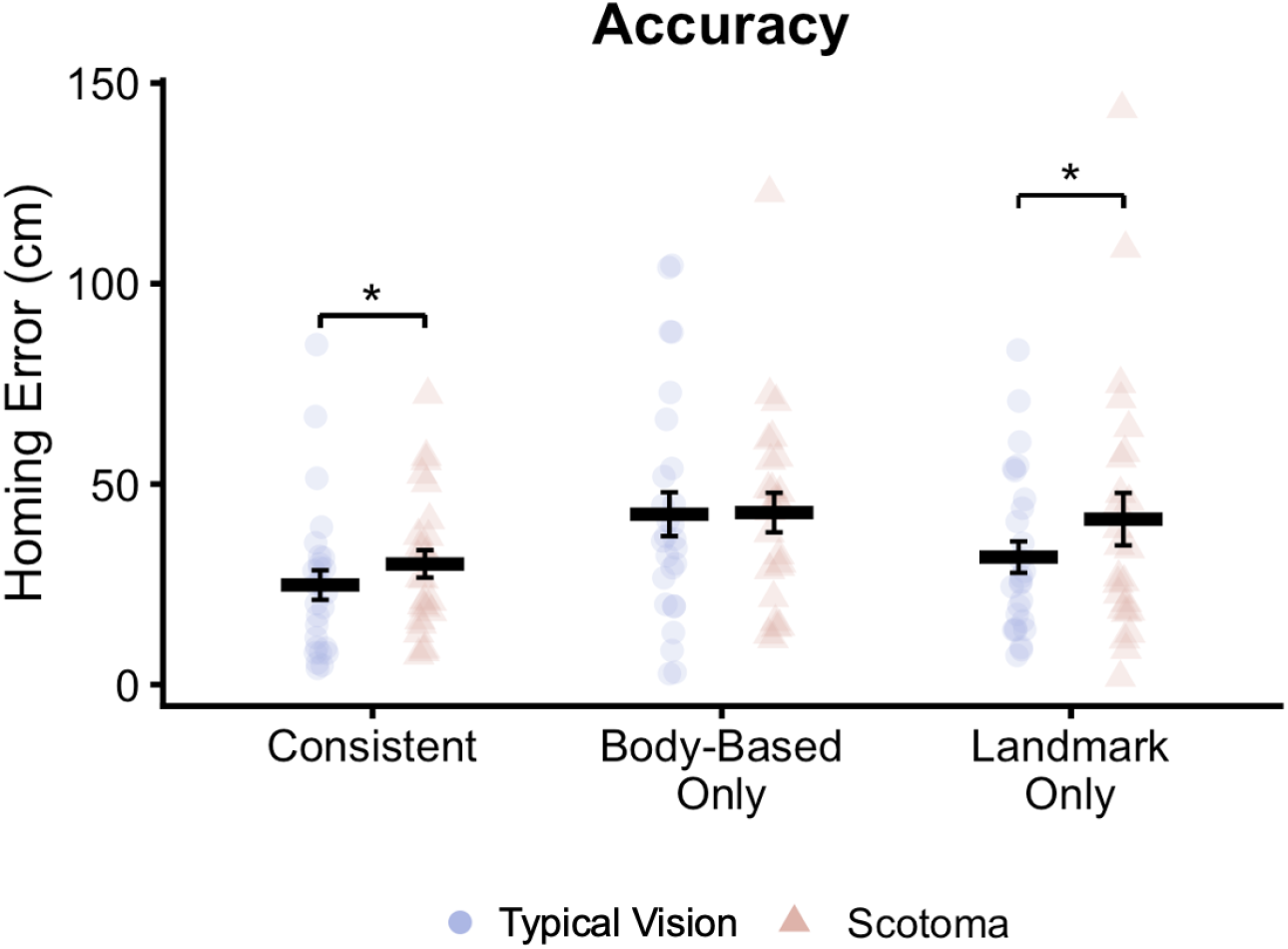
Homing error across cue conditions for typical vision (blue) and simulated scotoma (red). Points represent individual participant means and black horizontal lines represent posterior means with 95% credible intervals derived from the Bayesian multilevel models. Asterisks indicate conditions where the 95% credible interval for the difference between scotoma and typical vision excludes zero.

### Head Rotation

Trial-level yaw rotation rate was regressed onto cue condition, scotoma condition, and their interaction as fixed effects with a random intercept by participant to account for individual differences in baseline head rotation behavior. Yaw rotation rate in the consistent-cues condition under typical vision was 18.12^◦^/s (*B*_0_ = 18.12, 95% CrI [15.13, 21.10]; Figure 7). Yaw rotation rate was credibly greater in the body-based-only condition relative to consistent cues (*B* = 5.06, 95% CrI [3.08, 7.03], *BF*_10_ > 100), indicating that participants rotated their heads more rapidly when navigating without visual landmarks. The landmark-only condition did not credibly differ from consistent cues (*B* = 1.34, 95% CrI [*−*0.58, 3.30], *BF*_10_ = 0.49). Scotoma did not credibly affect yaw rotation rate in the consistent-cues condition (*B* = *−*0.36, 95% CrI [*−*2.31, 1.53], *BF*_10_ = 0.21). However, the interaction between scotoma and the body-based-only cue condition was credible (*B* = *−*3.17, 95% CrI [*−*5.87, *−*0.37], *BF*_10_ = 50), suggesting that the increased yaw rotation rate observed in the body-based-only condition under typical vision was reduced under simulated scotoma. The interaction between scotoma and landmark-only was not credible (*B* = *−*0.04, 95% CrI [*−*2.75, 2.67], *BF*_10_ = 0.22). Overall, participants increased head movements primarily when navigating without visual landmarks, consistent with greater exploratory behavior. The simulated central scotoma did not generally increase head scanning, suggesting that participants did not compensate for the scotoma by making larger head rotations during the task.

**Figure 7.**
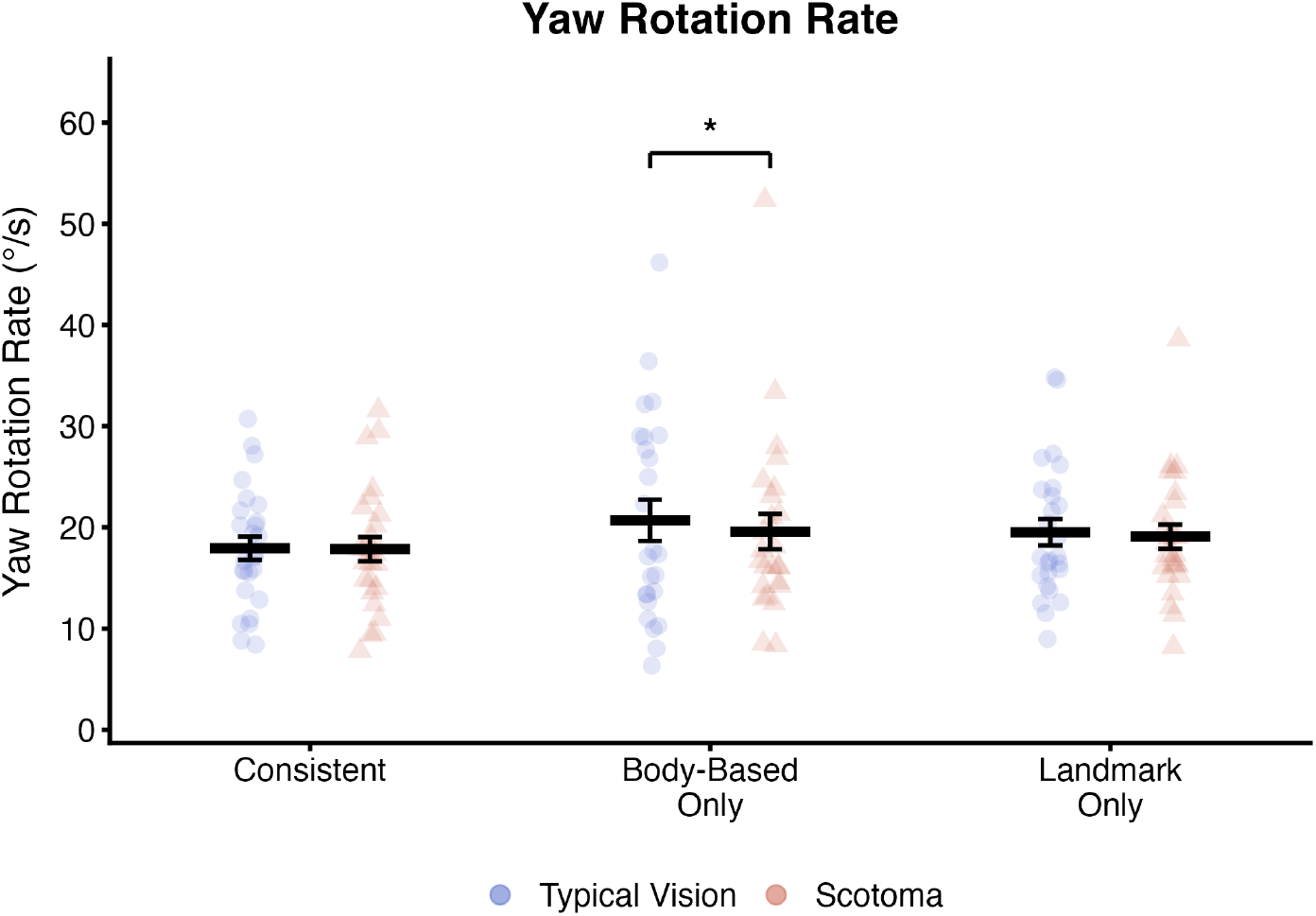
Mean yaw rotation rate (degrees per second) during the homing leg across cue conditions for typical vision (blue) and simulated scotoma (red). Points represent individual participant means and black horizontal lines represent group means with ±1 standard error. The asterisk indicates that the 95% credible interval for the interaction between scotoma and body-based cues excludes zero.

## Discussion

The present study examined how degrading central vision influences the use and integration of visual and body-based cues during spatial updating. The simulated scotoma occluded a large part of the viewer’s central visual field, thus reducing the availability of visual landmark information. We predicted that the reduced reliability of landmarks as a spatial cue would result in changes in performance in the scotoma condition showing 1) a smaller reduction in variability with two cues versus a single cue and 2) an increased weighting of body-based cues relative to visual landmarks, as compared to the typical vision condition. We also expected 3) error to increase when navigating with the scotoma, particularly in the conditions where landmarks were available.

For variability and cue weighting, we found overall similar patterns across scotoma and typical viewing conditions, contrary to our first two hypotheses. As expected based on prior work, variability was generally reduced when both cues were available compared to the single-cue conditions, but this reduction was smaller than that predicted by MLE in *both* typical and simulated scotoma conditions. Further, both conditions showed observed visual weights (derived from the conflict condition) that were higher than the predicted weights and did not differ from each other. Participants continued to overweight visual landmarks relative to predictions from optimal cue integration, regardless of vision condition, suggesting that acute central visual field loss reduced the reliability of visual information without altering the strategy for combining visual and body-based cues. As predicted in our third hypothesis, degraded central vision selectively increased homing error when visual landmarks were available but did not affect body-based homing. Along with the variability results, this suggests that central vision primarily contributes to the accuracy of visual landmark processing rather than to the precision of spatial updating. Finally, exploratory analyses provided little evidence that participants compensated for the simulated scotoma through increased head movements during navigation. Together, these findings suggest that acute central visual field loss degrades the quality of visual landmark information available for navigation, while the mechanisms governing multisensory cue integration remain remarkably robust.

The persistence of multisensory cue integration under simulated central visual field loss was reflected in the pattern of homing variability performance. Participants consistently benefited from having access to both visual and body-based information during spatial updating. Combining visual and body-based cues produced the clearest benefit for homing precision relative to landmark cues alone. While different from other work that often shows a greater benefit of two cues relative to body-based cues alone (Chen et al., 2017; Shayman, McCracken, Finney, Katsanevas, et al., 2024), our results may have been driven by the scotoma condition where body-based information could be used to reduce visual uncertainty (Chen et al., 2017). One explanation for the absence of a stronger multisensory benefit overall (as compared to the MLE prediction) is the substantial individual variability in cue weighting observed during spatial updating (Zanchi et al., 2022). Participants who naturally relied more heavily on visual landmarks may have gained relatively little from the addition of body-based information, whereas others may have benefited more from combining cues, reducing the overall group-level effect.

One of the most surprising findings was that simulated central visual field loss did not alter how participants combined visual and body-based information. Although participants consistently overweighted visual landmarks relative to predictions from MLE cue integration, this pattern was remarkably stable across both typical vision and simulated central visual field loss. The tendency to rely more heavily on visual information than predicted by MLE replicates previous work demonstrating that visual landmarks are often preferentially weighted during spatial updating under typical vision (Qi & Mou, 2024; Roy et al., 2023; Zhao & Warren, 2015b). More importantly, the absence of a shift in cue weighting suggests that acutely reducing the fidelity of central visual information was insufficient to trigger a change in participants’ cue-combination strategy. This finding is consistent with Shayman, McCracken, Finney, Fino, et al. (2024), who likewise found no change in cue weighting following simulated blur, but contrasts with findings suggesting that individuals with longstanding central visual field loss exhibit altered multisensory integration (Garcia et al., 2017). We purposefully designed this study as within-subjects so that we could directly compare cue weighting with and without vision loss within an individual, a benefit of VR simulation relative to studying those with clinical vision loss. However, factors related to participants experiencing both of the visual conditions (possible learning or order effects) and the implementation of a sudden change in vision need further study. Together, our results suggest that changes in cue weighting may depend less on the immediate degradation of visual information than on long-term perceptual adaptation. Participants in the present study experienced an acute simulated scotoma and therefore had little opportunity to recalibrate the relative reliability of visual and body-based information, whereas individuals with chronic central visual field loss have had extended experience adapting to altered sensory input.

Simulated central visual field loss selectively impaired homing performance accuracy when visual landmarks contributed to navigation, while leaving homing precision unaffected. This dissociation suggests that central vision contributes primarily to the fidelity of visual landmark information used during spatial updating rather than to the precision of the resulting spatial representation. In other words, degrading central vision increased systematic errors in landmark-based homing without making participants’ responses more variable from trial to trial. This interpretation is consistent with previous work suggesting that central visual field loss primarily impairs the acquisition of environmental information while leaving body-based self-motion processes relatively intact (Giudice, 2018). Likewise, previous homing studies have shown that the effectiveness of landmark information depends on its reliability, with greater accuracy observed when landmarks are more stable, located near the home location, or available in greater numbers (Qi & Mou, 2024; Roy et al., 2023; Zhao & Warren, 2015b). Together, these findings suggest that simulated central visual field loss reduced the fidelity, and therefore the reliability, of visual landmark information, leading to larger systematic homing errors when landmarks contributed to navigation while leaving the precision of spatial updating largely intact.

Exploratory head movements showed a different pattern than homing performance, suggesting that simulated central visual field loss influenced information-seeking behavior differently than homing accuracy. Participants reduced head rotation only when relying exclusively on body-based cues (likely due to the fact that moving the head would not help in this condition), whereas head movements remained unchanged when visual landmarks were available despite the degraded quality of those landmarks. One possible explanation is that participants continued to actively sample the environment when visual information could contribute to successful homing, even if central visual field loss reduced the fidelity of that information. In contrast, when visual landmarks were absent, exploratory head movements may have become less useful, leading participants to rely more directly on body-based information. This interpretation is speculative, though, because gaze behavior was not recorded. Future work combining eye and head tracking could determine whether navigators compensate for central visual field loss through altered visual sampling strategies or whether the observed changes reflect broader shifts in navigation behavior.

Several limitations should be considered when interpreting the present findings. First, the simulated scotoma was an opaque occlusion of fixed size and therefore did not capture several characteristics of clinical central visual field loss, including reduced acuity, reduced contrast sensitivity, perceptual filling-in, and the substantial variability in scotoma size observed across individuals with central visual field loss, such as macular degeneration (Hooper et al., 2008). We did not intend for the simulation to match a specific disease or stage of progression, but rather to demonstrate the broader consequences of reduced central visual field information. Ecologically valid simulations that more closely approximate the characteristics of clinical visual field loss will be important for determining how different aspects of loss may influence landmark use during spatial updating. Additionally, the scotoma was simulated in participants with typical vision, who consequently did not have the opportunity to develop the compensatory viewing strategies and perceptual adaptations that accompany long-standing central visual field loss. Future work should therefore examine cue combination during spatial updating in individuals with clinical central visual field loss to determine how long-term adaptation influences the use of visual and body-based cues.

## Conclusions & Implications

The present findings refine our understanding of how central vision contributes to human navigation by distinguishing its role in representing spatial information from its role in integrating sensory cues. Acute simulated central visual field loss selectively reduced the fidelity of visual landmark information, leading to less accurate landmark-based homing, yet it did not alter the precision of spatial updating or fundamentally change how visual and body-based cues were combined. These findings suggest that central vision is more critical for establishing accurate landmark representations than for the multisensory computations that integrate visual and body-based information during navigation.

This distinction has important implications for supporting navigation in individuals with central visual field loss. Rather than assuming a global impairment in spatial updating, interventions may be most effective when they enhance or supplement degraded landmark information while leveraging intact body-based navigation processes. Auditory and haptic landmarks, in particular, may provide valuable alternative sources of environ-mental information when visual landmarks are unreliable or unavailable (Giudice, 2018). Understanding how visual impairment changes navigation therefore requires distinguishing changes in the quality of sensory information from changes in the computations that combine that information.

Finally, our approach in this study used immersive VR and a gaze-contingent simulated scotoma which was possible because of advances in eye-tracking technologies within HMDs. VR provides an opportunity to implement controlled and ecologically valid studies on effects of visual information loss on navigation. While not a replacement for studies conducted with people with clinically-defined low vision, the approach provides a foundation for understanding effects of landmark degradation on navigation that could then be used to inform both cue-combination theory and application, including the development of navigation assistance and rehabilitation.

## Acknowledgments

The authors would like to thank the following individuals for help with data collection: Hollyn Gottschalk, CC Willemsen, Stevie Ayers, Ethan Butterfield, and Sultana Noffsinger.

This material is based upon work supported by the National Science Foundation Graduate Research Fellowship under Grant No. 2139322. Any opinions, findings, and conclusions or recommendations expressed in this material are those of the author(s) and do not necessarily reflect the views of the National Science Foundation.

Processed data and scripts are publicly available on the Open Science Framework (OSF): https://osf.io/vjg4x/overview?view_only=427b2c4bd15445a8af8f7731ae31d195. Experimental materials are available upon request to the corresponding author.

## Supplemental

### Bayesian Priors

Prior distributions were informed by Shayman, McCracken, Finney, Fino, et al. (2024), which examined homing performance under degraded visual conditions using the same virtual reality homing paradigm. Priors were chosen to reflect plausible parameter values observed in that study while remaining sufficiently broad to accommodate effects substantially larger than those previously reported. Prior predictive simulations were conducted to verify that the priors generated plausible values across the expected range of homing performance.

For the homing accuracy model, the intercept prior was specified as *N* (67, 25) to reflect expected baseline homing error. Fixed effects were assigned *N* (0, 30) priors, allowing large positive and negative condition effects while remaining centered on no effect. The residual standard deviation was assigned a *N* (40, 30) prior, and the participant-level random intercept standard deviation was assigned a weakly informative Exponential(0.05) prior.

Homing variability was modeled using a Gaussian distribution with an identity link. The intercept prior was specified as *N* (40, 20), reflecting an expected baseline variability of approximately 40 cm, consistent with Shayman, McCracken, Finney, Fino, et al. (2024). Fixed effects, including interaction terms, were assigned *N* (0, 15) priors, allowing substantial positive and negative changes in variability while remaining centered on no effect. The residual standard deviation was assigned an Exponential(0.05) prior.

Cue weighting was modeled using a Gaussian distribution with an identity link. The intercept prior was specified as *N* (0.5, 0.2), reflecting an expected baseline visual weight of 0.5, corresponding to equal weighting of visual and body-based cues. Fixed effects, including interaction terms, were assigned *N* (0, 0.2) priors, allowing variation in cue weighting while remaining centered on no effect. The residual standard deviation and participant-level random-intercept standard deviation were each assigned an Exponential(1) prior.

Mean yaw rotation rate was modeled using a Gaussian distribution with an identity link. The intercept prior was specified as *N* (18, 8), reflecting an expected baseline yaw rotation rate of approximately 18°/s while allowing substantial individual variability. Fixed effects, including interaction terms, were assigned *N* (0, 5) priors, allowing plausible positive and negative changes in rotation rate while remaining centered on no effect. The residual standard deviation and participant-level random-intercept standard deviation were each assigned an Exponential(0.1) prior.

## Footnotes

1 Because participants completed the typical vision and simulated scotoma conditions in counterbalanced order, supplementary analyses included session order as a fixed effect for all outcome variables. There were no credible main effects of session order for any outcome variable (all 95% CrIs included zero, all *BF*_10_ *<* 0.35), suggesting that the observed effects were not driven by session order.

